# Quantitative behavior of protein complexes in human cells

**DOI:** 10.1101/367227

**Authors:** Morteza H Chalabi, Vasileios Tsiamis, Lukas Käll, Fabio Vandin, Veit Schwämmle

**Author notes:** Biotech Research & Innovation Centre, University of Copenhagen, Copenhagen, Denmark (Morteza H Chalabi).

## Abstract

Translational and post-translational control mechanisms in the cell result in widely observable differences between measured gene transcription and protein abundances. Herein, protein complexes are among the most tightly controlled entities by selective degradation of their individual proteins. They furthermore act as control hubs that regulate highly important processes in the cell and exhibit a high functional diversity due to their ability to change their composition and their structure. To better understand and predict these functional states, extensive characterization of complex composition, behavior, and abundance is necessary. Mass spectrometry provides an unbiased approach to directly determine protein abundances across cell populations and thus to profile a comprehensive abundance map of proteins. We investigated the behavior of protein subunits in known complexes by comparing their abundance profiles across up to 140 cell types available in ProteomicsDB. After thorough assessment of different randomization methods and statistical scoring algorithms, we developed a computational tool to quantify the significance of concurrent profiles within a complex, therefore providing insights into the conservation of their composition across human cell types. We identified the intrinsic structures in complex behavior that allow to determine which proteins orchestrate complex function. This analysis can be extended to investigate common profiles within arbitrary protein groups. With the CoExpresso web service, we offer a potent scoring scheme to assess proteins for their co-regulation and thereby offer insight into their potential for forming functional groups like protein complexes. CoExpresso can be accessed through http://computproteomics/Apps/CoExpresso. Source code and R scripts for database generation are available at https://bitbucket.org/veitveit/coexpresso.

**Author summary:** Many proteins form multi-functional assemblies called protein complexes instead of working as singly units. These complexes control most processes in the cell making the full characterization of their behavior inevitable to understand cellular control mechanisms. Detailed knowledge about complex behavior will elucidate biomarkers and drug targets that exhibit and correct aberrant cell states, respectively. We investigated abundance changes of the protein complex components over more than 100 different human cell types. By using statistical scoring models, we estimated the evidence for the co-regulation of the proteins and revealed which proteins form subunits with impact on complex function and composition. By providing the interactive web service CoExpresso, any combination of proteins can be tested for their co-regulation in human cells.

## Introduction

Biological systems are governed by a multitude of entangled interactions between biomolecules with an immense number of physical and chemical properties. Protein complexes are large biomolecules with a wide range of tasks in the cell and consist of multiple subunits linked by non-covalent interactions. These interactions can lead to a variety of stable or transient states where the complexes display different compositions of their subunits or different structures that are often fine-tuned by post-translational modifications. An example of functional diversity are ribosomes that are known to contribute differentially to translation of distinct subpopulations of mRNAs [1]. There is a pressing need to investigate complex capabilities for regulatory control of cellular processes. To achieve this, a detailed map of protein complex composition, abundance, and behavior in different cell types and tissues is required. Such a map will considerably improve the characterization and the prediction of the functional states.

Various experimental methods exist to identify protein complexes and to determine and quantify which protein subunits they are composed of. Determination of protein interaction partners within a complex provides valuable knowledge about complex and protein function and thus their potential behavior [2]. Most prominent experimental methods to determine protein-protein interactions are based on the yeast-2-hybrid protocol or the application of affinity purification coupled with mass spectrometry [3,4]. These methods however suffer from either large false identification rates or depend on purification steps that often lead to a strong bias in the results. More details about protein structure can be achieved by chemical cross-linking or hydrogen-deuterium exchange mass spectrometry [5]. Despite the power of these methods, they cannot yet be applied on entire proteomes. For an accurate, large-scale and general characterization, protein complex behavior should be studied across large numbers of samples without perturbations towards e.g. subgroups of proteins and additionally rely on highly confident identification of the proteins.

There is an increasing amount of evidence supporting the hypothesis that the majority of protein complexes are tightly controlled in the cell. Post-transcriptional regulation occurs predominantly for protein complex members, leading to strong co-regulation of complex subunits. This could be shown by systematic investigation of protein and gene expression levels in human cancer [6,7], in a study comparing 11 cell types and 4 temporal states [8], based on the co-occurrence of protein pairs across human experiments in the PRIDE database [9], or generally in a selection of proteomics data sets [10]. In summary, these studies showed that only a fraction of complex composition and abundance is regulated at transcriptional level and therefore other mechanisms such as protein degradation contribute to protein complex stoichiometry. This highlights the power of directly measuring protein abundance profiles by common proteomics approaches such as bottom-up mass spectrometry to thoroughly study protein complexes and their variants across cell types and states.

In contrast to most proteomics data repositories where only raw data and identification results are available, ProteomicsDB [11,12] is a large compendium of quantitative protein abundances, therefore highly useful to investigate general patterns of protein changes across more than 100 different human cell lines.

Here, we apply three scoring models on the ProteomicsDB data to assess the significance of subunit co-regulation in protein complexes. We compare and benchmark different randomization and scoring approaches on known complexes and reveal particular substructures of complex behavior for a few selected use cases. The scoring and extensive visualization is implemented in the web service CoExpresso that allows investigating co-regulatory patterns in any group of human proteins.

**Fig 1.**
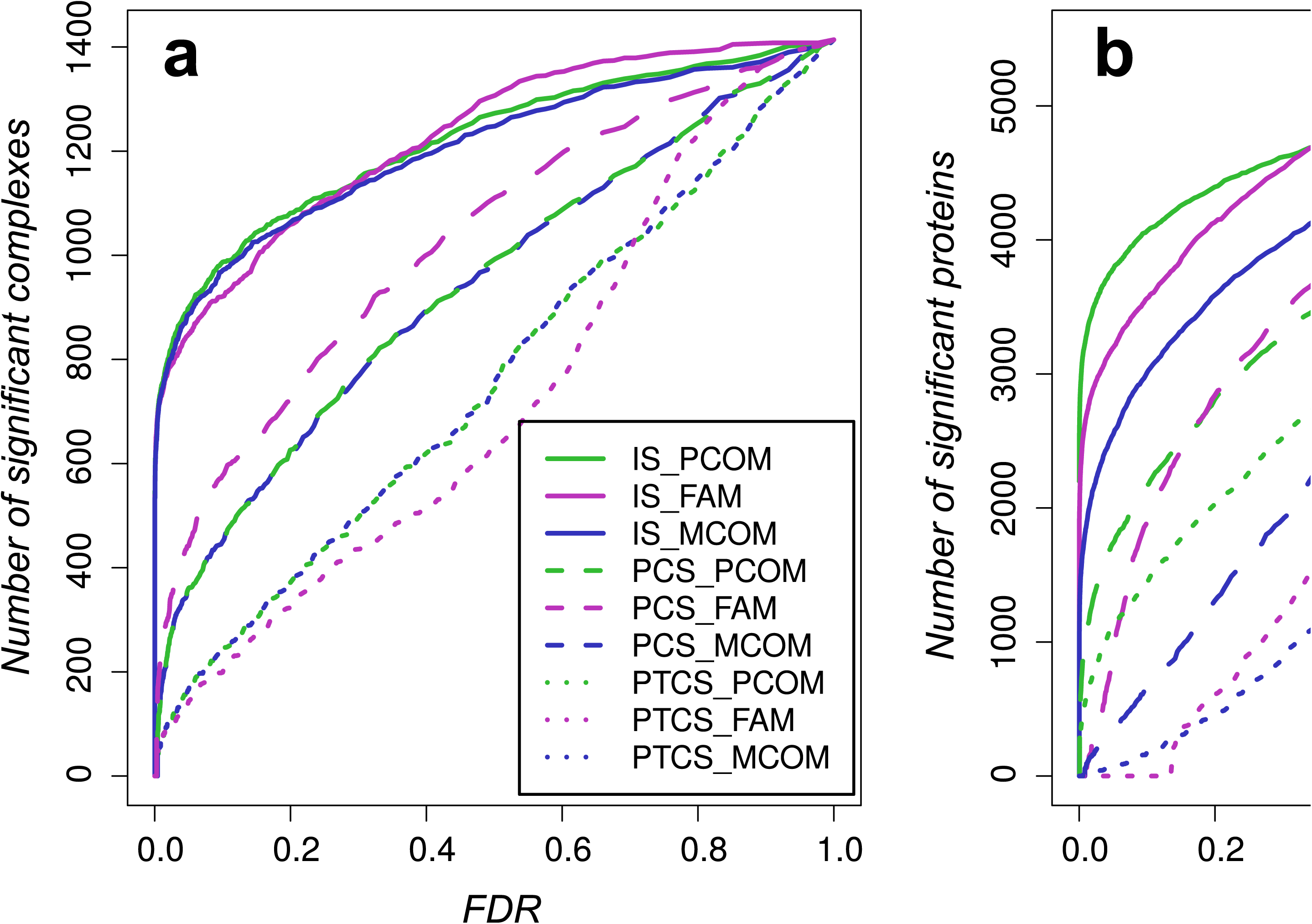
Schema of entire workflow to investigate protein complex behavior. Models and randomization methods that were not used in the final assessment of CORUM complexes are shown in grey. For a more detailed description of the workflow, see Methods.

## Results

We applied different scoring systems to evaluate whether proteins in human complexes exhibit similar regulatory behavior when compared over multiple cell types. Despite of having a large set of available protein abundances, coverage of the proteins over the 140 cell types was often sparse (S1 Fig), requiring scoring methods that account for missingness. Moreover, we needed to investigate different ways of randomizing the ProteomicsDB data to identify the best performing combination of scoring scheme and randomization procedure.

Table 1 summarizes the used methods and randomizations. In short, MCOM compares each protein profile versus the averaged profile of protein group, allowing to assess how much a protein follows this common trend. PCOM is based on pairwise comparisons and summarizes them by their sum. This method was implemented to consider internal structures of protein subgroups with high correlations. The FAM model is based on factor analysis and calculates weights for each protein, giving a measure of how much each protein contributes to the profile of the entire protein group.

For the following analysis, each protein complex reported in CORUM was assessed for coverage in ProteomicsDB and further evaluated by the different models when all protein subunits were available in at least 5 cell types. We tested a total of 1,414 protein groups annotated in CORUM.

**Table 1.**
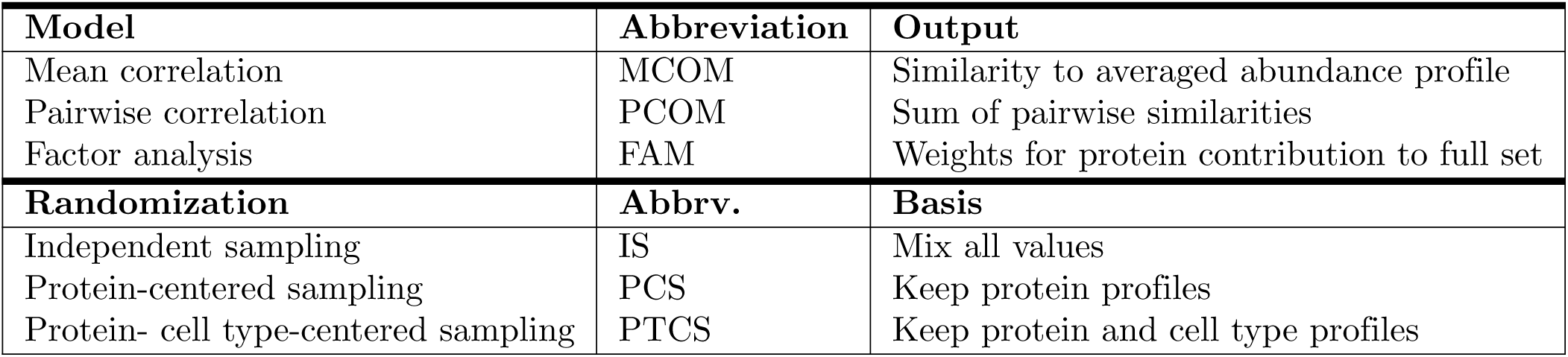
Summary of scoring models and randomization methods.

### Scoring models

By comparing the scores of the different models to scores obtained for randomly sampled data, we obtained probabilities to discard the observed abundance profiles as result of randomly chosen proteins. Thus the false discovery rates (FDRs), represented by p-values corrected for multiple testing, provide a measure for significance of a given complex on basis of co-regulation of its subunits within human cell types. The different randomization techniques were applied to resemble the intrinsic data structure on different scales.

Fig 2a compares the p-values calculated for each model and randomization. More “realistic” randomization (IS<PCS<PTCS) resulted in lower number of complexes with significant abundance profiles. MCOM and PCOM, both models being based on Pearson’s correlation, produced nearly the same results on complex level (see also S2 Fig). The FAMS approach however performed differently, reaching a higher number of significant complexes for the protein-centered randomization.

On protein level (Fig 2b), lower protein numbers with significant abundance profiles could be expected and were observed when using randomization methods that maintain protein and cell type properties. Here, PCOM displays a higher number of proteins than FAMS and MCOM for low false discovery rates.

**Fig 2.**
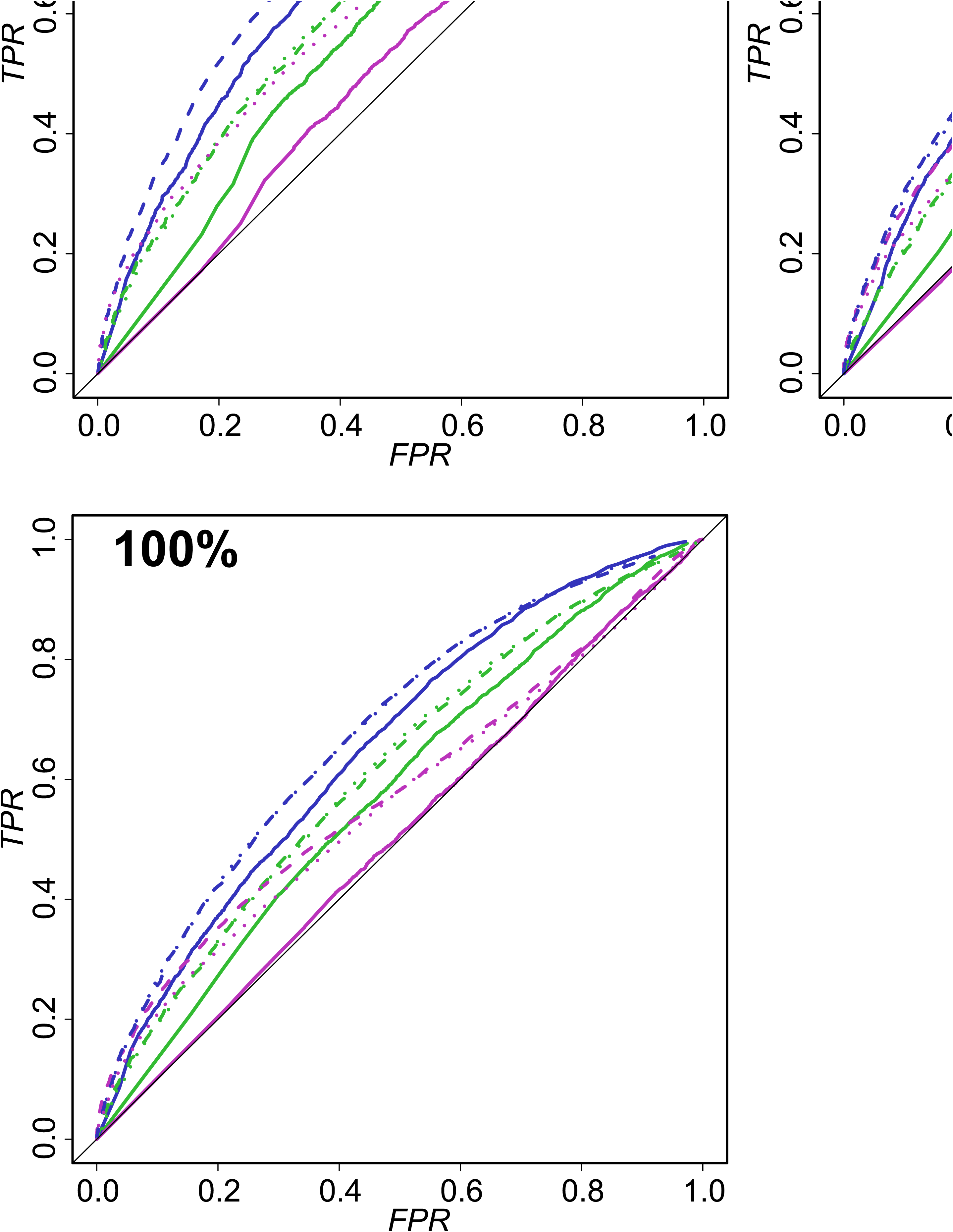
Comparison of models for significant co-regulations. Number of complexes (a) and proteins (b) with significant abundance profiles according to the different scoring models and randomizations calculated for different thresholds for their false discovery rate (FDR).

### Robustness

Recovery of proteins and complexes with significant abundance profiles does however not ensure robustness of the methods towards noise. As example, one could expect a protein complex to contain subunits that do not follow the general trend of the abundance profiles. This could be due to wrong assignment of a protein to a complex or due to different behavior of a subunit being heavily regulated by e.g. post-translational modifications or by forming transients regulating complex function.

Robustness to differently behaving proteins can be simulated by adding randomly chosen proteins to the CORUM complexes. In all complexes, we increased the number of proteins by 50%, 75% and 100%. Fig 3 shows ROC curves for these simulated complexes, where we compared the significance by counting true (actual complex subunits) and false positives (added proteins). Here, the different methods and randomization approaches showed consistent differences for their robustness. Randomization of the entire ProteomicsDB data lead to lower robustness for all methods. One the other hand, protein-centered (PCS) and protein-cell type centered (PTCS) randomization gave nearly identical performance results. Hence, the following analysis will focus on PCS randomization, although being the computationally mosts expensive one, as it yields higher counts of significant proteins. In addition, MCOM and FAM models had lower false positives rates at least in the lower range.

**Fig 3.**
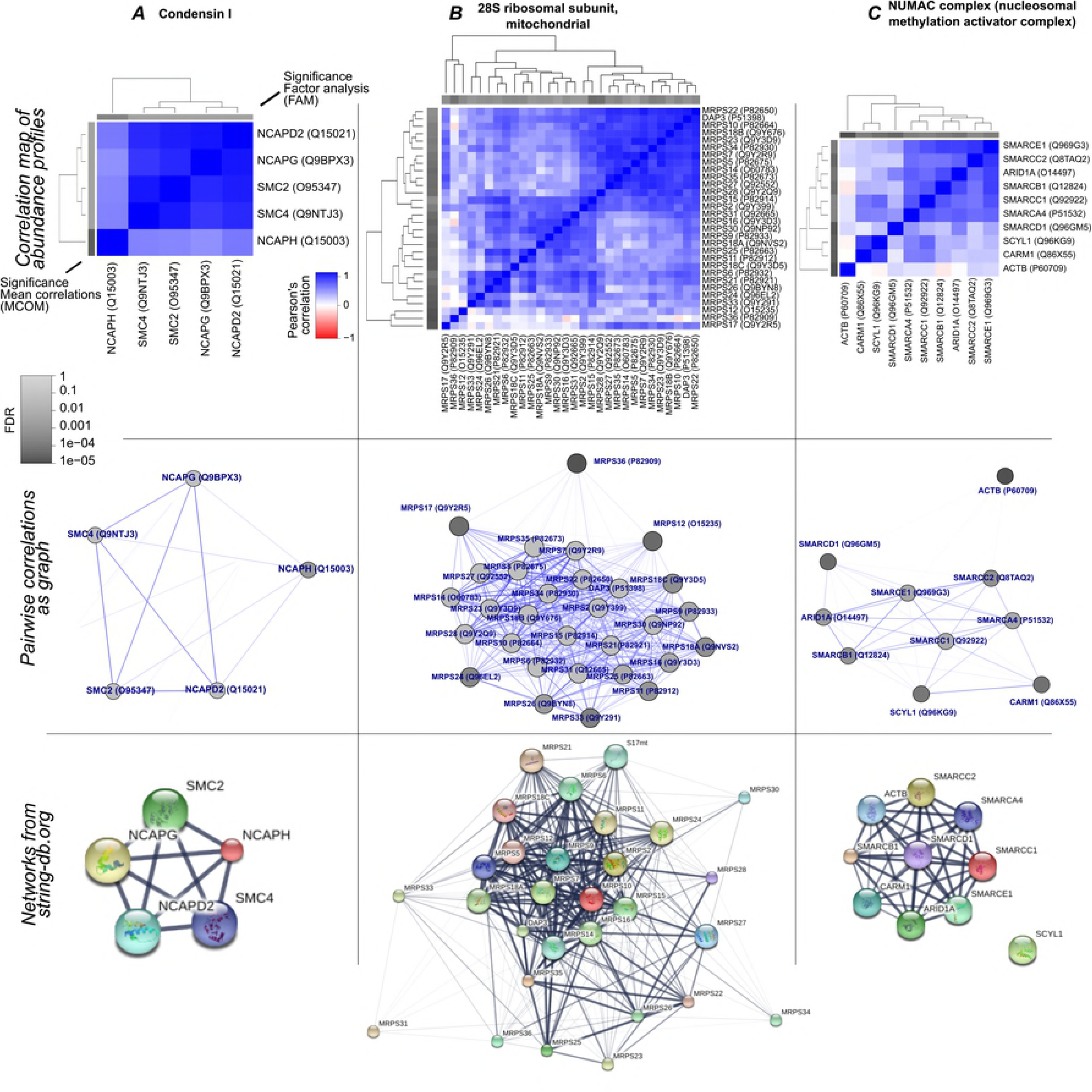
Performance of scoring models. measured by robustness to 50%, 75% and 100% artificially added random proteins. Proteins were categorized into complex subunits and random proteins. True positive and false positives rates (TPR and FPR) were given by the fraction of true positives and false positives at a given FDR threshold. MCOM and FAM models lead to better performance. Only slight difference between PCS and PTCS randomizations can be observed.

### Use cases

The following use cases will provide detailed results of the scoring models and general complex behavior for three selected complexes that are representative for the investigated complexes. We obtained 60 CORUM complexes with lowest FDR values (< 0.0003) for all three scoring models.

**Use case A**: Condensin I (Fig 4) represented the first of the complexes with lowest FDR values in all models (PCS randomization). All five proteins were commonly expressed in 75 cell types. Very high correlations between all proteins confirmed the high interaction evidence from STRINGdb [13]. However, Condensin subunit 2 (NCAPH) showed slightly lower correlation and lower scores. Indeed, NCAPH is known as regulatory subunit of Condensin I with different nucleolar localization during interphase [14]. We observed different abundance levels of NCAPH in several cell types leading to lower weight by factor analysis (S3 Fig).

**Use case B**: 28S mitochondrial ribosomal subunit (Fig 4), being essential for ATP production, represents the complex with lowest FDR in all models and most proteins. The 30 proteins were commonly available in 23 cell types. Both our visualization and STRINGdb interactions suggest a more open structure or composition of the complex with a core component of heavily co-regulated proteins. These proteins were mostly the same when comparing STRINGdb with our results and also were assigned higher significance by the here applied scoring models. Moreover, the correlation map (upper figure) roughly distinguishes 2 large subgroups with higher correlations amongst their proteins. We found a strikingly different behavior of these groups in lung tissue (S3 Fig). This suggests that the 28S ribosomal complex plays a different role in lung where it might break up into two functional units.

A group of the four proteins MRPS21, MRPS24, MRPS26 and MRPS33 exhibited highest correlations and reasonably high significance. We investigated their co-regulation as a protein group on their own where their abundance profiles were available in a higher number of 39 cell types and confirmed highly significant co-regulation for this higher coverage (not shown).

**Use case C**: NUMAC complex (nucleosomal methylation activator complex, Fig 4) denotes a case with slightly lower significance. All scoring models suggest high significance with an FDR below 0.5%. The 10 proteins were found in 33 cell types with ACTB distinct behavior and drastically higher abundance than the other proteins. Strong evidence for interactions of all components but SCYL1 in STRINGdb suggests that ACTB plays a crucial role in complex composition but might still have other functions in the cell. We assume that this protein is not actively degraded when not forming the complex. All 3 models agreed in having high FDR values for ACTB and SMARCD1 (FDR > 0.1), suggesting that the latter plays a particular role in this complex.

**Fig 4.**
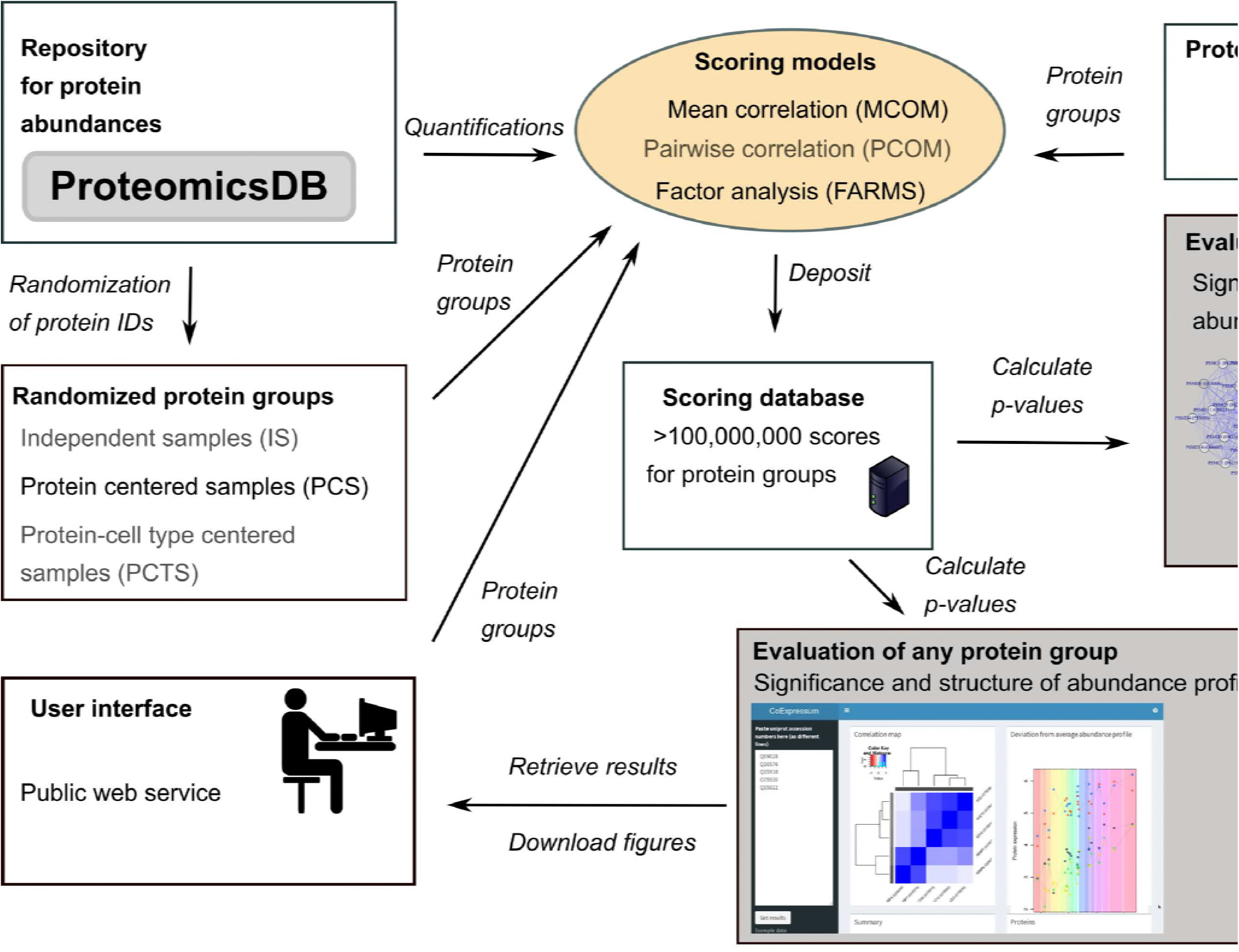
Examples for complexes with highly co-regulated proteins. Upper panels: hierarchical clustering of pairwise correlations between protein abundance profiles. The sidebars show the significance of MCOM and FAM models (PCS randomization). Middle panels: Network visualization of profile similarities. Edge widths correspond to pairwise correlations. Grey tones of the proteins depict FDR significance calculated by PCOM (PCS randomization). Lower panels: STRINGdb (version 10) networks of proteins. Edge width is given by interaction confidence.

### A data source for tightly co-regulated proteins

Given the strong co-regulation in annotated protein complexes, we asked whether our randomly sampled protein groups with highly significant co-regulation could determine novel but yet not well characterized complex compositions in human cells. Random protein groups with the highest scores did however not provide evidence for these proteins to be arranged as complexes as evidence scores for their interaction in STRINGdb were not sufficiently enriched. This does however not mean that highly significant protein groups do not have particular common biological functions such as co-regulation on transcriptional level or being common members of a pathway. We implemented CoExpresso that interrogates groups of human proteins to assess their co-regulation strength. The web service can be highly useful for the interested researcher to test their hypothesis on the basis of human cell types in general.

## Discussion

Published proteomics studies provide at least hundreds proteomics experiments per year from which a large percentage have their raw data deposited in the major data repositories (e.g. PRIDE nearly reaching 10,000 projects to date [15]). Availability of quantified protein abundances is however still very rare also because the comparison of protein abundance across experiments and projects is still a major bottleneck in the proteomics field. ProteomicsDB provides a large catalogue of protein abundances in human cell types which we used to thoroughly investigate protein complex behavior. Despite the large number of characterized cell types, data coverage is rather low, where more than 20% of the proteins were detected in only 2–5 cell types. Such low coverage hindered straight-forward application of e.g. simple correlation and we therefore compared a variety of different scoring models and randomizations that reproduce the inherent data structure.

Our comparison showed that appropriate randomizations are crucial to achieve results with simultaneously high recall and considerable robustness to the introduction of noise. The results speak against complete randomization of all values, where global differences amongst cell types and proteins are neglected. We found that protein identities (PCS method) needed to be maintained to reach robust results. On the other hand, maintaining the identity of the tissues (PCTS method) in the investigated protein group did not lead to lower robustness. We therefore conclude that testing properties of protein profiles in general should be compared to a randomized set where protein identity is kept. In data with many missing values, this randomization requires categorizing the random protein groups into their tissue coverage which can be computationally expensive. We therefore provide a web service that stores the randomizations and where arbitrary protein groups can be tested for their significance. By testing annotated complexes from the CORUM database for the significance of their concurrent protein abundance profiles, we could confirm almost 50% (500–600 depending on scoring model) of the protein groups being co-regulated with an FDR below 0.1. This confirms the tight regulation of complex proteins previously reported and extends this observation to be valid generally in human cells. Given the lack of coverage over sufficient cell types in many cases, resulting in rather low statistical power, we predict that most protein complexes will be found to be translationally and post-translationally regulated.

Our analysis additionally confirmed and extended details about protein complex substructure that indicates regulatory features that orchestrate complex function by changes in complex composition or by here not investigated post-translational modifications.

We furthermore tested whether the large database of randomized protein groups could be used to identify novel protein assemblies that represent highly interacting functional modules such as complexes. We did not find enrichment for known protein-protein interactions in the most significant protein groups. This means that investigating protein co-regulation by random sampling alone is not a good source to search for novel complexes but remains highly valuable to test for complex behavior and confirm their composition across cell types. Given the combinatorial explosion when considering the number of possible protein groupings, the random sampling strategy used here considers only a small fraction of all protein groups that contain highly co-regulated proteins. Novel protein assemblies could still be found by selective and iterative algorithms that determine protein groups with highest co-regulation within all possible combinations. The here presented study provides deep insight in to protein complex behavior in human cells. The data for all 1,414 investigated protein groups can be accessed via the CoExpresso web service. Arbitrary protein groups can be tested for their significance with respect to their co-regulation in human cell, such as investigating prior hypotheses about protein groups with common strongly co-regulated functional behavior. With more data on hand, we expect to improve statistical power and accuracy by including more data sets and by characterizing the role of quantified post-translational modifications.

## Materials and methods

Quantifications of proteins and IDs of known complexes were downloaded from ProteomicsDB [11, 12] and CORUM [16], respectively. We used three randomization approaches that differently resemble data structure within all protein abundance profiles. Scores were calculated for the co-regulation of proteins in a complex applying three different models for the comparison of protein profiles. The scores were stored in a database. For each protein in each complex, significance for their co-regulation was calculated and assessed on basis of the scores. A web service was implemented to allow interrogating the score database to test arbitrary protein groups for the significance of their co-regulation. Fig 1 provides an overview of the workflow and the web interface.

### Data retrieval

Quantitative abundance profiles of SwissProt proteins were extracted from ProteomicsDB hosting mass spectrometry based protein abundances for distinct human cell types including cell tissues, cell lines and fluids. In ProteomicsDB, proteins and samples are annotated according to UniProtKB and Brenda Tissue Ontology [17], respectively.

From the downloaded profiles (summer 2016), we retained only cell types with more than 1, 000 proteins and which were tagged by Brenda ontology terms. Proteins not available in at least 2 cell types were removed. This reduced the data to comprise 15,409 proteins and 140 tissues. Uniprot accession numbers for annotated human complexes were downloaded from CORUM and filtered for duplicates, leading to a total of 2175 reported complex compositions.

### Complex abundance profiles

For each protein group *C*, only the *n_t_* cell types with full coverage, *t* = [1..*n_t_*], i.e. having quantitative values for all proteins *p* = [1..*n_p_*], were considered, resulting in a *n_t_* by *n_p_* matrix *E_C_*(*t,p*).

#### Randomization techniques

We applied 3 different forms of randomization to obtain random protein groups being quantified in the same number of cell types as the proteins of protein group *C*. The often relatively low coverage of proteins over multiple cell types required creating randomized sets for each combination of number of cell types and number of proteins.

*Independent sampling (IS):* Randomization of quantitative values of all proteins in all tissues comprised sets with the same dimensions as the to be tested protein group. That is, the *n_t_* by *n_p_* randomized values were obtained by sampling, independently at random, *n_t_n_p_* values from all quantitative values of all proteins in all tissues. 10,000 random groups were created for each combination of *n_t_* and *n_p_*.

*Protein-centered sampling (PCS):* Randomization of proteins and categorization into cell type coverage. This randomization type turned out to be more complex and a sufficient large coverage of random groups was achieved by the following procedure. For each combination of number of proteins *n_p_* and number of cell types *n_t_*:

1. Take all proteins being each quantified in at least *n_t_* cell types
2. Repeat the following 5, 000 times: sample *n_p_* proteins IDs and count full cell type coverage of the protein group
3. Keep unique protein combinations with coverage over at least five cell types

With this procedure, we obtained 1,000-20,000 unique and random protein groups for each relevant combination giving a total of more than 20,000,000 randomized groups.

*Protein- and tissue-centered sampling (PTCS):* All proteins simultaneously found in the same cell types as the tested protein group were randomized to create 10,000 samples. That is, *n_p_* proteins are sampled independently at random from all proteins that appear in the same cell types as the tested protein group, and their observed values in those cell types are considered.

### Similarity models and scoring

*Mean correlation model (MCOM):* Protein abundances were averaged for each cell type, restricting to cell types covered by the entire protein group, *M*(*t*) =< *E_C_*(*t,p*) >*_p_*. For each protein *p*, Pearson’s correlation to the means *M*(*t*) provides a measure of how much the protein follows the common profile of the protein group,

*S*_MCOM_(*p*) = cor(*M*(*t*), *E_C_*(*t,p*)), where cor(*x*(*t*), *y*(*t*)) denotes Pearson’s correlation between samples *x* and *y*.

*Pairwise correlation model (PCOM):* Pearson’s correlation was calculated between all proteins pairs using the abundances in the cell types covered by all proteins. The score is then given by the sum,

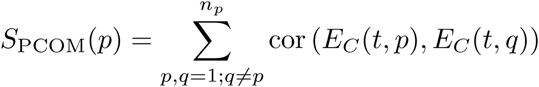

*Factor analysis model (FAMS):* The model is based on factor analysis developed for microarray analysis [18] and recently modified to improve protein inference in bottom-up mass spectrometry data [19]. The following parameters were used: Weight *w* = 0.1, *μ* = 0.1, 1,000 maximal iterations and a minimal noise of 0.0001. The feature weights *W* were used to score each protein of a group: *S*_FARMS_(*p*) = *W*(*p*).

*Scores for protein groups:* Overall scores per protein group were generated by simply averaging the scores of the individual proteins, 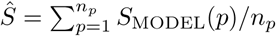, where MODEL stands for either MCOM, PCOM or FARMS.

### Scoring statistics

For each model, randomization method, and a given combination of *n_t_* and *n_p_*, the aforementioned scores were calculated for randomized protein groups, and stored in a database. These scores, more than 100,000,000 in total, were then used to calculate the probabilities to reject the null hypothesis (of observing the score for a set of *n_p_* proteins over *n_t_* tissues) for both a single protein *p* and a group of proteins:

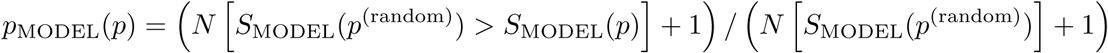
and

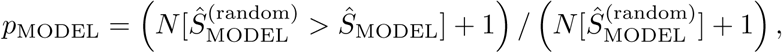
where *p*^(random)^ denotes a protein from a randomized protein group, 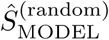 a score for a randomized protein group, and *N*[…] counting the number of all valid cases within the brackets.

For *p*-values from multiple protein groups, correction for multiple testing was carried out via the Benjamini-Hochberg procedure.

## Supporting information

**S1 Fig. Low coverage of protein profiles across cell types complicates co-expression analysis.**

**S2 Fig. Comparison of complex p-values between models. Colors indicate cell type coverage.**

**S3 Fig. Protein abundance over cell types for 3 protein complexes.** Left panels: Abundance is shown versus the mean taken for each cell type (different colors). Middle panels: Protein abundances for the different cell types which were ordered according to the mean expression. Line thickness corresponds to the weights calculated by the FAM model.

